# C_4_-like *Sesuvium sesuvioides* (Aizoaceae) exhibits CAM in cotyledons and putative C_4_+CAM metabolism in adult leaves as revealed by transcriptome analysis

**DOI:** 10.1101/2023.09.29.560146

**Authors:** Christian Siadjeu, Gudrun Kadereit

## Abstract

The co-occurrence of C_4_ and CAM photosynthesis in a single species seems to be unusual and rare, probably because of the difficulty to co-regulate both pathways effectively. Nevertheless, it represents a unique chance in gaining new insights into the evolution and regulation of these complex pathways. Comparative transcriptomics using RNA-seq revealed C_4_-like and CAM photosynthesis in *Sesuvium sesuvioides* (Aizoaceae) leaves and cotyledons, respectively. When compared to cotyledons, phosphoenolpyruvate carboxylase 4 (PEPC4) and some key C_4_ genes were found to be up regulated in leaves. During the day, the expression of NADP-dependent malic enzyme (NADP-ME) was significantly higher in cotyledons than in leaves. The acidity titration confirmed higher acidity in the morning than in the previous evening indicating the induction of weak CAM in cotyledons by environmental conditions. Comparison of the leaves of *S. sesuvioides* (C_4_-like) and *S. portulacastrum* (C_3_) revealed that PEPC1 was significantly higher in *S. sesuvioides*, while PEPC3 and PEPC4 were up-regulated in *S. portulacastrum*. Finally, potential key regulatory elements involved in the C_4_ and CAM pathways were identified. These findings provide a new species in which C_4_ and CAM co-occur and raises the question if this phenomenon is indeed so rare or just hard to detect and probably more common in succulent C_4_ lineages.

**Highlight:** C_4_ and CAM metabolism co-occur in the C_4_-like species *Sesuvium sesuvioides* (Aizoaceae).

## Introduction

In a number of eudicot families, C_4_ photosynthesis evolved in ancestrally succulent lineages (see Berasategui et al., submitted, for an overview), prominent examples are Chenopodiaceae (Kadereit et al., 2012), Aizoaceae-Sesuvioideae (Bohley et al., 2015), Portulacaceae (Ocampo & Columbus, 2012) and Zygophyllaceae (Bellstedt et al., 2012). For some of these families also Crassulacean Acid Metabolism (CAM) is known or even the predominant carbon concentrating mechanism (CCM), such as Aizoaceae and Portulacaceae. Both CCMs share the same core metabolic enzymes, both evolved repeatedly multiple times, however, it seems that they rarely co-occur. So far, the co-occurrence of C_4_ and CAM was verified for only four genera, *Portulaca* (Portulacaceae, Koch and Kennedy, 1980), *Spinifex* (Poaceae; Ho et al., 2019), *Ottelia* (Hydrocharitaceae; Han et al., 2020) and *Trianthema* (Aizoaceae; Winter et al., 2021). Recent evidence demonstrated that C_4_ and CAM are operating in the same cells in *Portulaca oleracea* under drought conditions (Moreno-Villena et al., 2022). This integration is plausible due to several copies of core C_4_ genes (i.e. PEPC) that are recruited for C_4_ and CAM, respectively, sharing a set of biochemical reactions. Detecting the co-occurrence of C_4_ and CAM is tedious, requires living collections and an experimental approach, which is why this phenomenon has not been documented very often. However, we hypothesize that it might be more common in succulent C_4_ lineages than currently known.

C_4_ photosynthesis is an adaptive evolutionary response to the harmful effect of photorespiration under hot and dry growing conditions, by concentrating CO_2_ around RuBisCO (Sage, 2003). This allows for a remarkably efficient photosynthesis, as well as water and nitrogen use. The C_4_ pathway is a complex combination of anatomical and biochemical specialization. In succulent C_4_ lineages the C_4_ anatomy is particularly diverse (Berasategui et al., submitted; Voznesenskaya et al., 2010). Often the Kranz cells are not arranged as an inner wreath around the vascular bundles like in plants with typical Kranz anatomy, but form a continuous inner chlorenchyma layer around the central water storage tissue of the leaf (e.g., Schüssler et al., 2017; Bohley et al., 2015). The general pathway in which CO_2_ is converted to bicarbonate (HCO_3_^-^) by carbonic anhydrase in mesophyll (M) cells and then fixed to the 3-carbon molecule phospho*enol*pyruvate by the enzyme Phosphoenolpyruvate Carboxylase (PEPC) to form the 4-carbon molecule oxaloacetate unifies all plants with C_4_ photosynthesis. Oxaloacetate is then either reduced to malate or transaminated to aspartate. After diffusing to an adjacent Kranz cell, malate or aspartate is primarily decarboxylated by the enzymes either NADP-dependent malic enzyme (NADP-ME) or NAD-dependent malic enzyme (NAD-ME). This decarboxylation releases CO_2_ in high concentrations around RuBisCO and ensures high photosynthetic efficiency. This carbon concentration mechanism (CCM) is supported and facilitated by a decrease in the ratio of M to Kranz cells as opposed to the C_3_ ancestors. Unlike C_4_ photosynthesis, the CCM of CAM photosynthesis is temporally asynchronous in a single-cell system. During the night, plants open stomata, and CO_2_ is fixed and converted to malate, which is stored in the vacuole as malic acid. During the day, stomata are closed and stored malate is transported out the vacuole and decarboxylated to release CO_2_ that is then fixed by RuBisCO and enters the Calvin cycle for sugar production. This asynchronous carbon fixation system allows plants to keep their stomata closed to avoid water loss through evapotranspiration during the hottest period of the day. Thus, plants with this type of metabolism are able to grow in hot and dry environments. While it is straight forward to detect obligate CAM plants by means of a strong carbon isotope signal and consistent differences between morning and evening acid concentrations, it is laborious to detect facultative or weak CAM plants that only induce CAM under stress (Messerschmid et al., 2021; Winter, 2019). Weak CAM can neither be detected by carbon isotope ratios in C_3_ species nor in C_4_ species. In C_3_ species, the discrimination of RuBisCO towards the heavier C isotope and in C_4_ species the much higher activity of the PEPC in the C_4_ pathway subdue the low CAM signal.

Aizoaceae comprise annual or perennial herbs, rarely shrubs or trees growing in tropical and subtropical regions, predominantly in South Africa (Hartmann, 2017). Most species of the family are succulent and many, especially from subfamily *Mesembryanthemoideae* and *Ruschioideae* are documented CAM plants (Messerschmid et al., 2021). In Aizoaceae, C_4_ photosynthesis is restricted to subfamily Sesuvioideae and likely evolved multiple times (Bohley et al., 2015). A striking diversity of leaf anatomical types and the occurrence of both biochemical subtypes of C_4_ (NAD-ME and NADP-ME) can be observed. In addition to this photosynthetic diversity, two species from Sesuvioideae have been reported to activate low CAM under drought, i.e., the C_3_ species *Sesuvium portulacastrum* and the C_4_ species *Trianthema portulacastrum* (Winter et al., 2019; Winter et al., 2021). Yet another species of this subfamily arouses curiosity: *Sesuvium sesuvioides*, a succulent C_4_-like species with uncommon C_4_ features and photosynthetic plasticity during leaf aging. Structural, physiological, and biochemical analysis of *Sesuvium sesuvioides* indicated a relatively high mesophyll-bundle sheath cell ratio and the presence of RuBisCO large subunit together with PEPC in the mesophyll cells (Bohley et al., 2019). Furthermore, a decrease of C_4_ enzyme activities was observed from young to mature to senescent leaves (Bohley et al., 2019). Although Bohley et al. (2019) did not observe any CAM activity under well-watered conditions, they did not eliminate the existence of CAM under dry conditions.

Such species that exhibit photosynthetic variability may contain footprints left from the evolution of CAM and C_4_ photosynthesis and thus provide useful information to either disentangle or gain new insight into the evolution and regulation of C_4_ and CAM metabolism. From this perspective, the photosynthetic flexibility in subfamily *Sesuvioideae* represents an excellent potential model. We need such models as with climate warming, many agricultural regions approaching their potential peak of productivity, and with an estimated population of 10 billion people by 2050, C_4_ and CAM represent a promising way to increase productivity and hence yield to meet global demands for food owing to their intrinsic ability to adapt to hot and dry environments. Thus, both CCMs are targets for genetic engineering into C_3_ species. Substantial efforts taken in the past to introduce C_4_ and CAM features into C_3_ plants failed to reach the envisioned goals due to lack of knowledge of C_4_ and CAM photosynthesis at the system level. Therefore, mechanisms underlying C_4_ and CAM anatomical structure, gene-specific expression, and regulation network in C_4_ must be clarified further (Cui, 2021), and each new mosaic stone will help to solve the conundrum. To test our hypothesis that *S. sesuvioides* operates combined C_4_ and CAM photosynthesis, we (1) performed a comparative transcriptome analysis between cotyledons and young leaves of *S. sesuvioides* (C_4_-like species) and young leaves of *Sesuvium portulacastrum* (C_3_ species) under stressful conditions: high light intensity and drought. We then (2) investigated the integration of C_4_+CAM in *S. sesuvioides* via the identification of candidate genes linked to C_4_ and CAM previously identified in *Portulaca* (Christin et al., 2020). Finally, we explored the regulatory elements controlling C_4_ and C_3_ pathways. Our analyses revealed that *S. sesuvioides* is operating weak CAM in cotyledons and C_4_+CAM in leaves as proved by gene expression analysis and supported by acid titration. Moreover, C_4_+CAM candidate genes were found up-regulated during the day suggesting the integration of C_4_+CAM metabolism in *S. sesuvioides*.

## Materials and Methods

### Plant materials

Plants of *S. sesuvioides* (C_4_-like) and *S. portulacastrum* (C_3_) were grown from seeds and cuttings respectively, in the experimental greenhouse of the Munich-Nymphenburg Botanical Garden, Germany. *Sesuvium sesuvioides* seeds were collected from a location situated ∼80 km east of Sendelingsdrif, Karas, Namibia [∼28.20946°S, 17.28936°E, 208 m altitude, voucher: Klak 2431 (BOL)] and *S. portulacastrum* plant materials were collected in Texas, USA (MSB Serial number 0394523; year collected: 2007). For simplicity, we will sometimes use C_4_ instead of C_4_-like. Germinated seedlings (about 1cm) and one-year-old plants of *S. sesuvioides* and one-year-old plants of *S. portulacastrum* were transferred to climate chambers with the following parameters: photoperiod light/dark 14h/10h, 60% humidity, [CO_2_] = 400ppm, maximum light intensity =785 μmol/m^2^/s, day/night temperature of 25/22°C. Plants were watered every 2 days. This unexpectedly created stressful conditions specifically drought with changes in leaves colours as shown in the pictures (Supplementary Fig. S1). Samples for transcriptome were harvested two weeks after transferring the plants to the climate chamber. Three young adult leaves of three plants of each species were collected and for *S. sesuvioides* cotyledons three replicates of four plants were harvested during day and night.

### RNA extraction and sequencing

Total RNA was extracted from leaves and cotyledons as described by Siadjeu et al. (2021) using innuPREP Plant RNA Kit (Analytik Jena AG, Jena, Germany). Total RNA quality control was performed using the 2100 Bioanalyzer (Agilent Technologies) and Agarose gel electrophoresis. Messenger RNA was purified from total RNA using poly-T oligo-attached magnetic beads. After quality control and fragmentation, the first-strand cDNA was synthesized using random hexamer primers followed by the second-strand cDNA synthesis. The library was ready after end repair, A-tailing, adapter ligation, size selection, amplification, and purification.

### Transcriptome analysis

Sequence read quality control was assessed using FastQC (https://www.bioinformatics.babraham.ac.uk/projects/fastqc/) and summarized with MultiQC (Ewels et al., 2016). Random sequencing errors in reads were corrected with a k-mer-based method implemented in Rcorrector (Song and Florea, 2015) and uncorrectable reads were removed from the reads using TranscriptomeAssemblyTools (Fig. 1) (https://github.com/harvardinformatics/TranscriptomeAssemblyTools). Low-quality reads and adapters were filtered using TrimGalore v0.6.7 (https://github.com/FelixKrueger/TrimGalore/releases). The rRNA reads were removed by aligning trimmed reads against the SILVA v-138 rRNA database using Bowtie2 v.2.4.5 (Langmead and Salzberg, 2012). De novo transcriptome assembly was performed using Trinity v.2.14.0 (Grabherr et al., 2013) with the following parameters (Trinity --seqType fq -- SS_lib_type RF --max_memory 200G --min_contig_length 300 --CPU 16). The de novo transcriptome assembly quality was first confirmed by aligning cleaned reads back to the corresponding de novo transcriptome assembly using Bowtie2 v.2.4.5. Secondly, the BUSCO score (odb10) was determined using BUSCO v.4 (Manni et al., 2021). For the downstream analysis, the initial assembly of each species was processed as follows. We reduced the transcriptome data by clustering transcripts with 98% similarity with CD-HIT v4.8.1 (Fu et al., 2012). Then, we selected only transcripts harboring coding sequences with TransDecoder v.5.5.0 (https://github.com/sghignone/TransDecoder). TransDecoder performs a precomputed blastX alignment to the Uniprot protein sequence database to improve the prediction of open reading frames. Finally, we used the indexes of transcripts containing coding sequences to subset the initial assemblies. The reduced transcriptomes were used for differential expression analysis. For differential expression analysis, we exclusively focused on *S. sesuvioides* when comparing leaves collected during the day to cotyledons collected both during the day and at night. For the comparison between C_3_ and C_4_-like species, we used young adult leaves collected during the day.

**Fig. 1:**
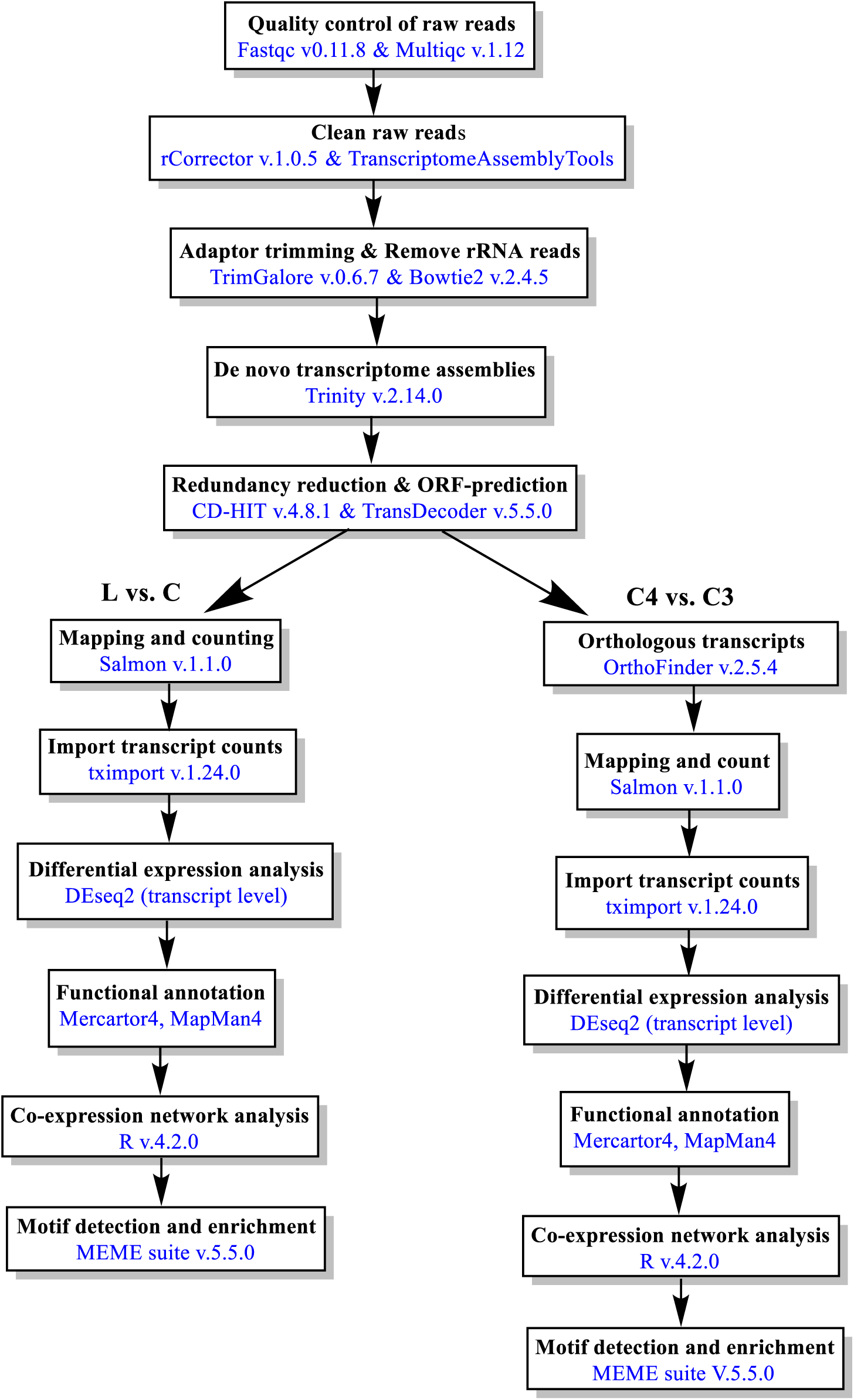
Workflow of transcriptome data analysis. The blue color represents the software used. L: leaves, C: cotyledons

### Transcript quantification and differential expression analysis

We quantified transcript abundance with Salmon by aligning the reads of each species to its reduced transcriptome (Fig. 1). The software tximport v. 1.24.0 (Soneson et al., 2016) was used to import transcript level abundances, estimated counts, and effective lengths for differential expression analysis. For comparison between the leaves of C_3_ and C_4_-like species, we searched for orthologs between their reduced transcriptomes using OrthoFinder v2.5.4 (Emms and Kelly, 2019). Based on the orthology between C_3_ and C_4_-like species, we obtained unique indexes by blasting C_3_ and C_4_ transcripts against each other. The best blast hits with lowest e-value and high bit scores were selected and considered homologous. The C_4_ transcript was used as a reference for the unique Id. If a C_4_ transcript was hit multiple times with the transcript of a C_3_ species, only one transcript was kept to get a similar number of transcripts for differential expression analysis. Finally, the unique C_4_ Id was changed in the Salmon output of the C_3_ species before importing the data with tximport (Script is available at https://github.com/Siadjeu/Sesuvioideae_C4-CAM). We assessed differential expression with the program Deseq2 (Love et al., 2014). Transcripts with p-value and p-adjusted as false discovery rate < 0.05 were considered significantly expressed andLog2FC was set > 1.

### Pathway and gene ontology (GO) annotation

Metabolic pathways and annotations of differentially expressed transcripts (DETs) were assigned via the tool Mercator4 (Schwacke et al., 2019). Swiss-prot protein sequences database and prot-scriber were included to improve the annotations. Unassigned transcripts were manually assigned based on the knowledge of the molecular functions in C_4_/CAM, photorespiration, and starch metabolism. We assigned GO terms to DETs using Blast2GO through OmicsBox with cutoff = 55, GO weight =5, e-value = 1.e-5, HSP-hit coverage cutoff = 80 and hit filter =500. We enriched the GO terms using Fisher’s exact test via OmicsBox.

### Co-expression network analysis

The co-expression analysis was performed using the unsupervised machine learning algorithm k-means in R. We used the three most popular methods for determining the optimal cluster: the Elbow and silhouette (Rousseeuw, 1987) methods and gap statistic (Tibshirnai et al., 2001). The normalized read counts of DETs were used. The maximum number of clusters (k) was set to 10. If transcription factors (TFs) and phytohormones were clustered with C_4_ or CAM genes, they were considered candidate TFs and phytohormones controlling C_4_ or CAM photosynthesis. The k-means clustering script is available underhttps://github.com/Siadjeu/Sesuvioideae_C4-CAM.

### Motif detection and enrichment

Transcript sequences of k-means clusters were analyzed for motif identification and enrichment. The program MEME suite v.5.5.0 (Bailey et al., 2015) was deployed to detect de novo motifs with the following parameters: E-value threshold =0.05, minimum motif size = 6 bp. We checked for motif redundancy with TOMTOM (Gupta et al., 2007) using the motif database JASPAR nonredundant core 2022. We enriched the detected motifs using AME (Mcleay and Bailey, 2010) with the following parameters:ame --verbose 1 --oc. --scoring avg --method fisher --hit-lo-fraction 0.25 --evalue-report-threshold 0.05 --control --shuffle kmer 2 MemeUpC_3_vsC_4_photoStach.fasta motif_db/JASPAR.

### Titratable acidity

Putative CAM activity under stressful conditions was investigated in cotyledons and leaves of *S. sesuvioides* and leaves of *S. portulacastrum* via comparative titratable acidity between 30 minutes before the end and 30 minutes before the beginning of the light period (19.30h and 5.30h, respectively). Cotyledons and leaves were harvested and snap-frozen in liquid nitrogen and stored at - 20°C. Since cotyledons were small, eight plants of *S. sesuvioides* for each harvesting time were collected. Stored cotyledons and leaves were chopped and weighed. About 50mg of the cotyledons and leaves material were incubated at 60 °C in 20% ethanol for 60 minutes. The extract obtained was aliquoted into three replicates of the same volume (1 mL). The extracted acid was neutralized by adding 0.01 M NaOH in 1ul increments (Silvera et al., 2005).

## Results

### Transcription assembly and quality assessment

The initial transcriptome assemblies of *S. sesuvioides* and *S. portulacastrum* contained 313,669 and 248,314 transcripts that were highly complete and less fragmented (C:96.2%, F:2.1) and (C:95.9 F:2.4), respectively (Table 1). Only 27% of transcripts (85,170) of *S. sesuvioides* and 31% of *S. portulacastrum* (76,264) were predicted to possess coding sequences. However, no significant changes were observed in the BUSCO scores (*S. sesuvioides*: C:95.3%, F:2.5%; *S. portulacastrum*: C:95.1%, F:2.8%). Transcripts with coding sequences were used for differential expression analysis between cotyledons and leaves of *S. sesuvioides*. For cross-species differential expression analysis, 43,769 and 40,429 orthologous transcripts were identified in *S. sesuvioides* (C:84.9%, F:3.5%) and *S. portulacastrum* (C:82.8%, F:4.6%) with an overall alignment of 73% and 76%, respectively.

**Table 1:** Transcriptome assemblies of *Sesuvium sesuvioides* and *S. portulacastrum*.

| Species | Software | Number of<br>contig | % of reads<br>alignment | Complete<br>BUSCO (%) | Fragmented<br>BUSCO (%) |
| --- | --- | --- | --- | --- | --- |
| *Sesuvium sesuvioides* (C<sub>4</sub>) | Trinity | 313669 | 98.79 | 96.2 | 2.1 |
|  | CD-HIT | 232335 | 98.76 | 95.5 | 2.5 |
|  | TransDecoder | 85170 | 93.25 | 95.3 | 2.5 |
|  | OrthoFinder | 43769 | 73.05 | 84.9 | 3.5 |
| *Sesuvium portulacastrum* (C<sub>3</sub>) | Trinity | 248314 | 98.62 | 95.9 | 2.4 |
|  | CD-HIT | 195534 | 98.57 | 95.4 | 2.8 |
|  | TransDecoder | 76264 | 90.65 | 95.1 | 2.8 |
|  | OrthoFinder | 40429 | 75.7 | 82.8 | 4.6 |

### Differential expression analysis across *Sesuvium* species

Two comparative analyses were carried out: (1) between leaves and cotyledons of *S. sesuvioides*, and (2) between leaves of *S. sesuvioides* and leaves of *S. portulacastrum*. Leaves and cotyledons of *S. sesuvioides* were clearly separated based on their expression profile (Fig. 2a). Likewise, C_3_ and C_4_ species were clustered according to their photosynthetic type mainly along PC1 (Fig. 2b). A total of 6,063 transcripts were found to be significantly differentially expressed between leaves and cotyledons of *S. sesuvioides* during the day (L and CD), of which 2,492 were up-regulated in leaves and 3,571 in cotyledons (Fig. 2c). When comparing leaves during the day (L) and cotyledons of *S. sesuvioides* during the night (CN), 1242 were up-regulated in leaves and 2,941 in cotyledons. Comparison of cotyledons between night (CN) and day (CD) revealed that 1,706 and 821 transcripts were found to be up-regulated at night and day, respectively. Between the C_3_ (*S. portulacastrum*) and C_4_ (*S. sesuvioides*) species, we found 20,867 orthologous transcripts in a 1:1 relationship. Out of these orthologs, 3,860 transcripts were significantly up-regulated in the C_3_ species and 2,433 in the C_4_-like species (Fig. 2c).

**Fig. 2:**
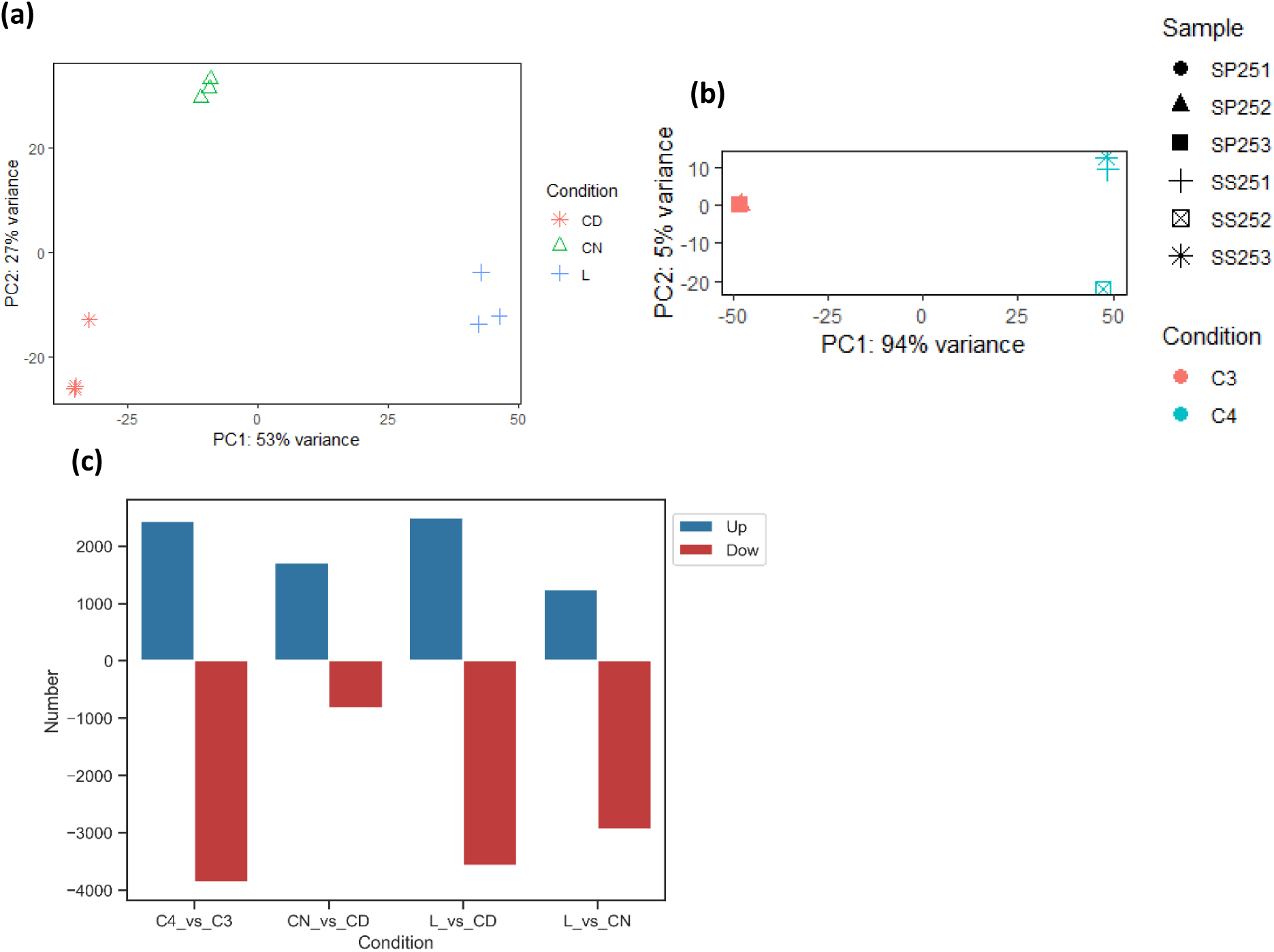
Transcriptome expression patterns of *Sesuvium*. (a) Principal component analysis (PCA) of leaves vs cotyledons of *S. sesuvioides*, CD: cotyledons day, CN cotyledons night, L: leaves. (b) PCA of *S. sesuvioides* vs *S. portulacastrum*, SP251:253: *S. portulacastrum* replicates 1:3, SS251: *S. sesuvioides* replicates 1:3. (c) Number of differential expressed transcripts: C4_vs_C3, between leaves of *S. sesuvioides* (C_4_-like) and *S. portulacastrum* (C_3_); CN_vs_CD, between cotyledons of *S. sesuvioides* collected during the night and during the day; L_vs_CD, between leaves and cotyledons of *S. sesuvioides* both collected during the day; L_vs_CD, between leaves and cotyledons of *S. sesuvioides* both collected during the night.

### Functional annotation of differentially expressed transcripts

To analyze photosynthesis in leaves and cotyledons of *S. sesuvioides*, we identified differentially expressed transcripts (p < 0.05, log2FC > 1) related to CCM according to annotation (Fig. 3). The transcripts related to CCM were subsequently clustered based on their role in decarboxylation, Calvin cycle, carboxylation, citrate generation, PEPC regeneration and regulation, photorespiration, transfer acid generation, and transporter. We then counted the number of related transcripts of each cluster to identify the photosynthetic mode of cotyledons and leaves (Fig. 3a). We found that transcripts involved in Calvin cycle, photorespiration, carboxylation, and decarboxylation were abundant in cotyledons as compared to leaves, whereas transporters involved in CCMs (C_4_ and CAM), transfer acid generation and PEPC regeneration were higher in leaves than in cotyledons (especially those collected during the day). This suggests that different CCMs are acting in leaves and cotyledons of *S. sesuvioides*. To investigate further, we analyzed specific genes related to the functional categories. In both CCMs, CO_2_ is fixed and converted to HCO_3_^-^ enzymatically by a beta-carbonic anhydrase (CA) in mesophyll cells (Gowik and Westhoff, 2011; Winter and Smith, 2022). HCO_3_^-^ is then fixed to the 3-carbon molecule phospho*enol*pyruvate (PEP) by the enzyme PEPC into the 4-carbon molecule oxaloacetate. Oxaloacetate is then reduced to either malate by NAD(P)-malate dehydrogenase (NAD(P)-MDH or transaminated by aspartate aminotransferase (AST) to aspartate. In this process, the enzyme phosphoenolpyruvate carboxylase kinase 1 (PPCK1) ensures the regulation of PEPC. After diffusing to the adjacent Kranz cell layer, malate or aspartate is decarboxylated and high concentrations of CO_2_ accumulate around RuBisCO allowing high rates of carboxylation. However, CAM differs from C_4_ by a nocturnal CO_2_ fixation and accumulation of malate or citrate in the vacuole, and a diurnal decarboxylation of accumulated malate by the malic enzymes (e.g. NADP-ME). Our results showed that transcripts encoding carboxylation enzymes (PEPC4 and PPCK1) were upregulated in leaves as compared to cotyledons during the day, while transcripts encoding decarboxylation enzymes NADP-ME and chloroplastic NADP-ME4 were significantly abundant in cotyledons at night (Fig. 4a). These results indicate a higher decarboxylation rate in cotyledons during the day and night. Thus, we suspected putative CAM and C_4_ photosynthesis in cotyledons and leaves of *Sesuvium sesuvioides*, respectively. To determine whether CAM is occurring in the cotyledons, we then compared the transcript profiles of CD and CN (Fig. 3a).

**Fig. 3:**
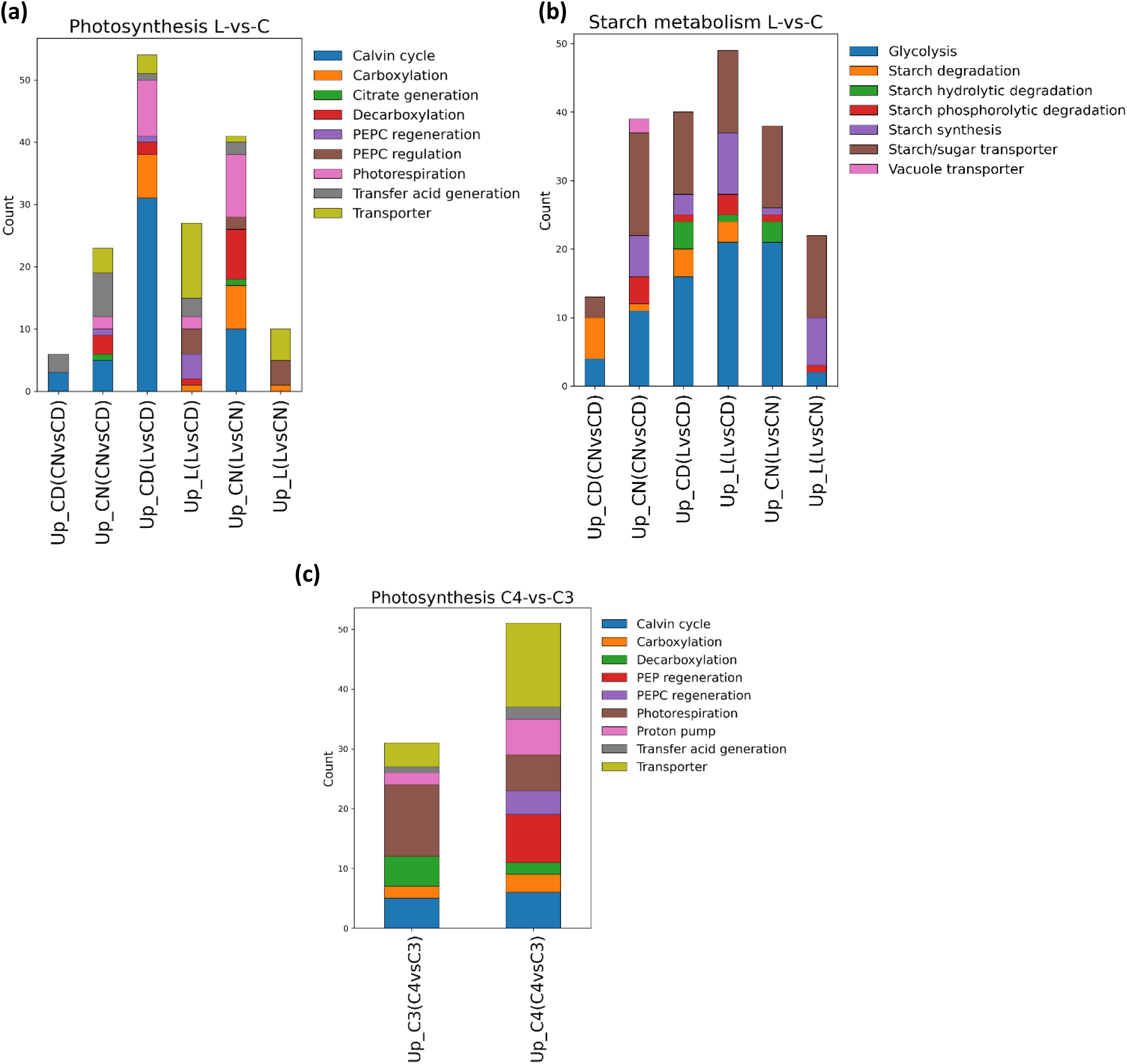
Stacked bar charts of functional annotated differentially expressed genes and abundance of selected genes involved in CCM and starch metabolism. (a) Functional annotated differentially expressed genes related to photosynthesis between leaves (L) and cotyledons (C) of *Sesuvium sesuvioides*. (b) Functional annotated differentially expressed genes related to starch metabolism between L and C. (c) Functional annotated differential expressed genes related to photosynthesis between C_3_ and C_4_ species

**Fig. 4.**
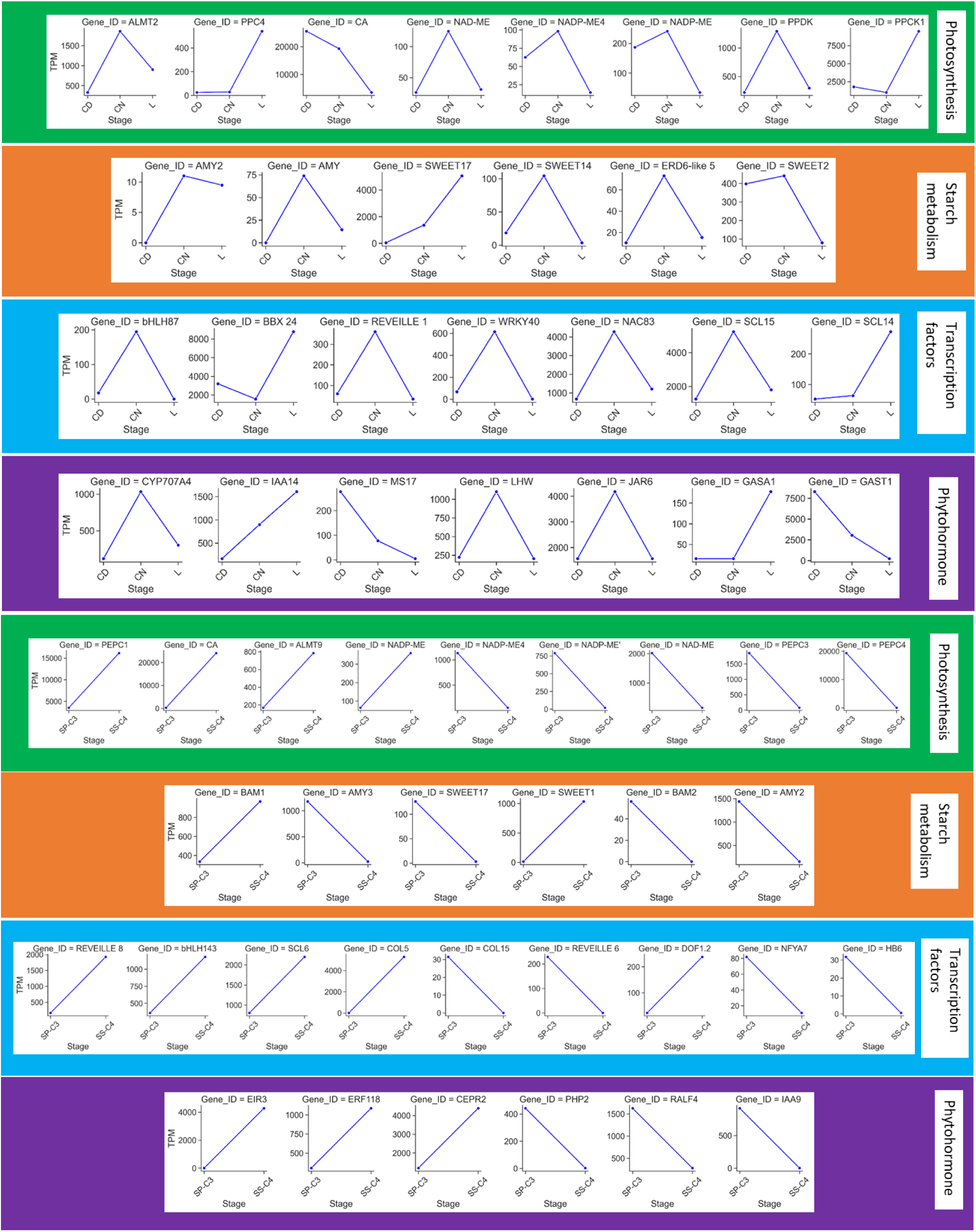
Abundance of selected genes classified in functional categories. (a):(d) abundance between leaves and cotyledons. (e):(h) abundance between C_3_ and C_4_ species. BBX: B-box, bZIP: Basic Leucine zipper, CEPR: C-terminally Encoded Peptide Receptor, COL: CONSTANS-Like, CYP707A4: cytochrome P450, EIR: Ethylene insensitive root, ERF: Ethylene-responsive element binding factor, GARP: Golden2, ARR-B, Psr1, GASA: gibberellic acid-stimulated Arabidopsis, HB: Homeobox, IAA: indole-3-acetic acid, JAR: JASMONATE RESISTENT, LHW: LONESOME HIGHWAY, NAC (NAM, ATAF and CUC), NFYA: Nuclear transcription factor Y subunit alpha, PHP: Pseudo histidine phosphotransfer protein, SCL: SCARECROW-LIKE, SWEET: Sugar Will Eventually be Exported Transporters

In CAM plants, malate generated at night is supposedly transported to the vacuole by a malate transporter Aluminium-activated malate transporter (ALMT) (Borland et al., 2009). We observed a significant increase in transcripts of ALMT2 in CN (Fig. 4a). The vacuolar malate influx is driven by the difference in membrane potential established by vacuolar-type proton adenosine triphosphatase ATPase (VHA) and the pyrophosphate-energized membrane proton pump (AVP) (Smith et al., 1996). However, as in *Portulaca* (Moreno-Villena et al., 2022), no significant expression of these genes was observed in the CN. Although no significant expression of decarboxylation enzymes NADP-ME in cotyledons during the night when compared to CD, a mitochondrial NAD-ME was highly abundant in CN (Fig. 4a). However, we found that RuBisCo small unit (RbcS1) (Supplementary Dataset S1-S2) was up-regulated in CD. This suggested that the decarboxylation activity of NAD-ME at night is aimed primarily at malate respiration (Tronconi et al., 2008). Additionally, ATP-citrate synthase alpha chain protein 3 (ACL-3) which is involved in citrate synthesis (Winter and Smith, 2022) was found up-regulated in CN. Citrate is produced when acetyl CoA reacts with oxaloacetate suggesting that citrate contributes as well to the acidification in *S. sesuvioides* at night.

Another feature of CAM photosynthesis is the nightly regeneration of PEP via phosphorolytic starch degradation (Heyduk et al., 2019). Moreover, Moreno-Villena et al. (2022) hypothesized that CAM induction is associated with the increase in sugar transporter. The number of transcripts related to starch/sugar transporter and starch phosphorolytic degradation was high in CN (Fig. 3b). We found that genes the probable alpha-amylase 2 (AMY2), alpha-amylase (AMY) and alpha-1,4 glucan phosphorylase L-2 isozyme (PHO2) involved in starch phosphorolytic degradation were significantly abundant at night (Fig. 4b). The sugar transporters, SWEET14, SWEET17, ERD6-like 5 were up-regulated at night (Fig. 4b). It is worth mentioning that we did not observe a significant expression of PEPC at night. However, pyruvate phosphate dikinase (PPDK) was significantly abundant in the cotyledons at night (Fig. 4a). The enzyme PPDK catalyzes the regeneration of the CO_2_ acceptor PEP via pyruvate (Chastain et al., 2011). Taking all this molecular evidence, our data suggest weak CAM induction in cotyledons, which was further confirmed by titratable acidity. Indeed, we found a significant accumulation of acids overnight (t test, adjusted p-value=0.018) (Fig. 5). The acidity difference between the morning and night corresponds to weak CAM induction. Thus, *S. sesuvioides* is a C_4_-like species operating weak CAM photosynthesis in cotyledons.

**Fig. 5.**
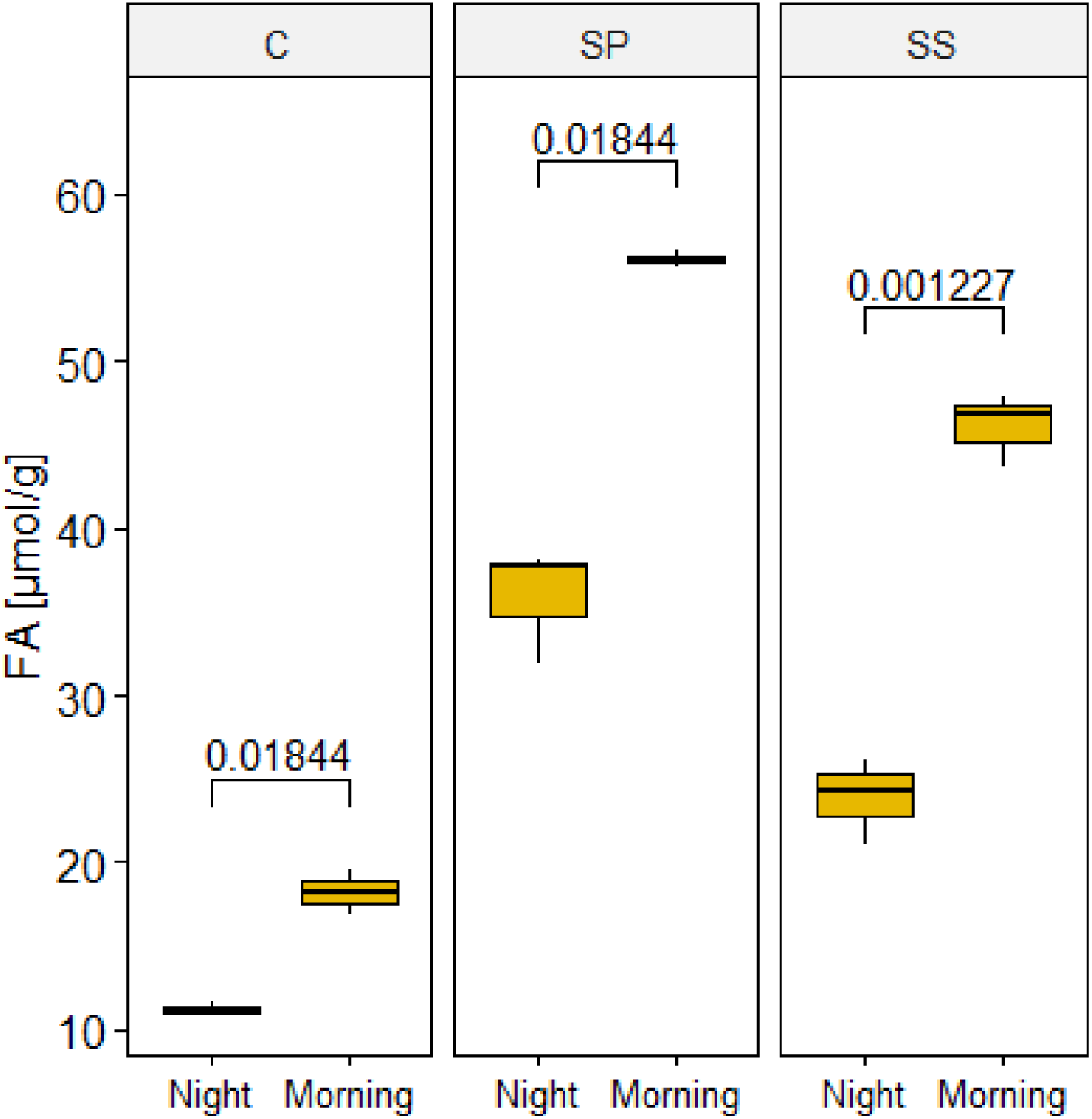
Boxplots of morning-night differences in titratable acidity of cotyledons (C) and leaves (SS) of *S. sesuvioides* and leaves of *S. portulacastrum* (SP). Values in the graph indicate the adjusted p values of significant differences between morning and night

To explore the difference between ancestral C_3_ photosynthesis to C_4_-like photosynthesis in *S. sesuvioides*, we compared the expression profile of *S. portulacastrum* (C_3_) and *S. sesuvioides* (C_4_-like). Differentially expressed transcripts between these species were clustered according to photosynthetic sub-pathways. We found a significant accumulation of genes involved in C_4_-related pathways (except PEP and PEPC regenerations that were only found in the C_4_-like species) in both species (Fig. 3c). The number of genes involved in carboxylation, proton pump, transfer acid generation, and transporter were higher in *S. sesuvioides* (C_4_-like; Fig. 3c). Conversely, genes related to decarboxylation and photorespiration were abundant in the C_3_ species as compared to C_4_-like species.

We then investigated genes related to carboxylation and decarboxylation. Surprisingly, while PEPC1 was up-regulated in the C_4_-like species, PEPC3 and PEPC4 were significantly expressed in the C_3_ species (Fig. 4e). Pyruvate phosphate dikinase (PPDK), and PPCK1 were up-regulated in the C_4_-like species (Supplementary Dataset S3). The decarboxylation enzymes chloroplastic NADP-ME4, NADP-ME, and NAD-ME were significantly accumulated in the C_3_ species. However, another NADP-ME copy was up-regulated in the C_4_-like species. These findings suggest that *S. sesuvioides* as C_4_-like species employs NADP-ME as a decarboxylation enzyme but can additionally use NAD-ME (Supplementary Dataset S3). The findings also suggest that during our experiment CAM photosynthesis was induced in *S. portulacastrum*. It was shown indeed that *S. portulacastrun* is capable of inducing CAM in stressful conditions (Ting, 1989, Winter et al., 2019). Gene ontology (GO) enrichment showed that response to stress was among the top 20 categories that were enriched in both species (Fig. 6). Moreover, we found that ALMT9 and ALMT4 were significantly up-regulated in *S. sesuvioides* and *S. portulacastrum,* respectively (Fig. 4e, Supplementary Dataset S3-S4). ALMT was originally responsible for nocturnal malate accumulation caused by an inward-rectifying anion-selective channel that forces only malate influx to the vacuole (Hafke et al., 2003). However, Meyer et al. (2011) demonstrated that AtALMT6 functions as a malate influx or efflux channel depending on the tonoplast potential. Several copies of transcripts of genes that control the tonoplast potential were significantly up-regulated in *S. sesuvioides* (VHA-A, VHA-C, VHA-E1) and in *S. portulacastrum* (VHA-G1, VHA-B1) (Fig. 7). This indicates diurnal vacuolar malate efflux in both species, hence the possibility of weak CAM being induced in *S. sesuvioides* and *S. portulacastrum*.

**Fig. 6.**
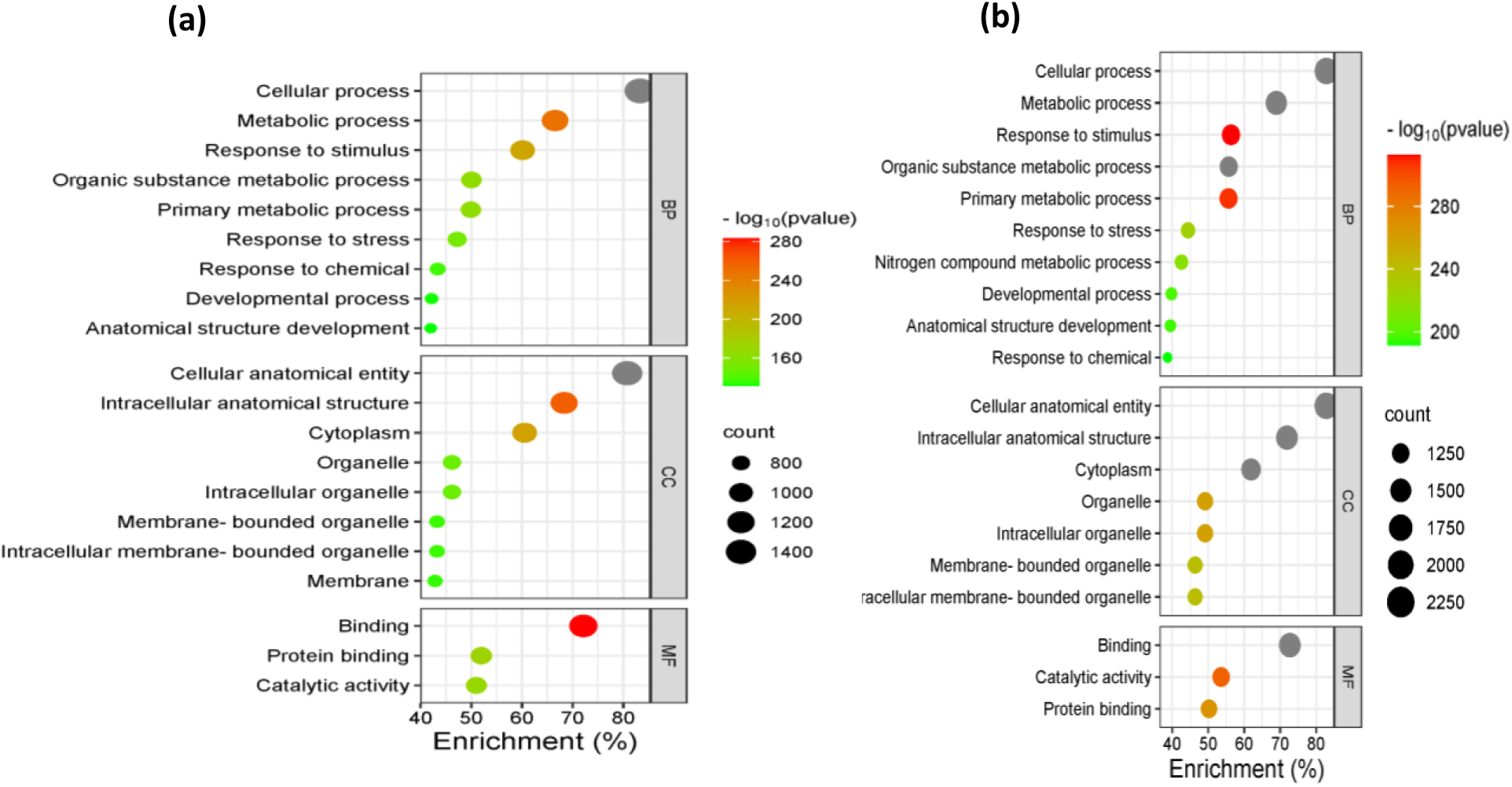
Gene ontology enrichment. (a) *S. sesuvioides*, (b) *S. portulacastrum*

**Fig. 7.**
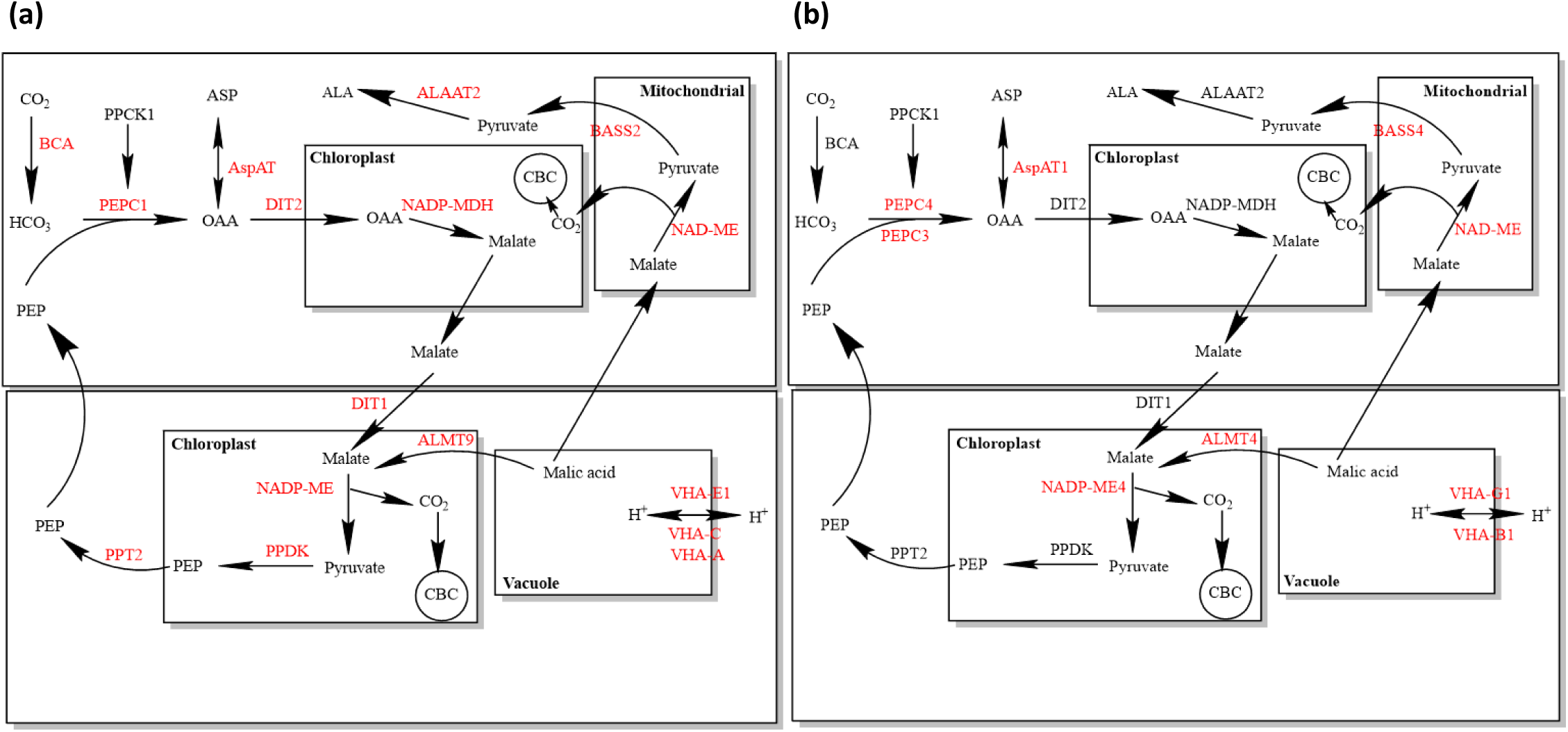
Putative photosynthetic cycle in *S. sesuvioides* (a) and *S. portulacastrum* (b). Red colour stands for genes that were up-regulated

### Regulation and hormonal signaling in *Sesuvium*

To identify regulation and signaling elements, we performed an unsupervised k-means clustering. The three methods used showed the best k was two (Supplementary Fig. S2). We found that almost all transcripts involved in photosynthesis, starch metabolism, transcription factor, and phytohormone signaling were grouped together (Cluster with the highest number of transcripts) for all comparisons (Fig. 8, Supplementary Dataset S5). To identify the most abundant TF families controlling regulation, we clustered TFs found in groups including photosynthesis, starch metabolism, and phytohormone signaling according to their families (Fig. 9).

**Fig. 8.**
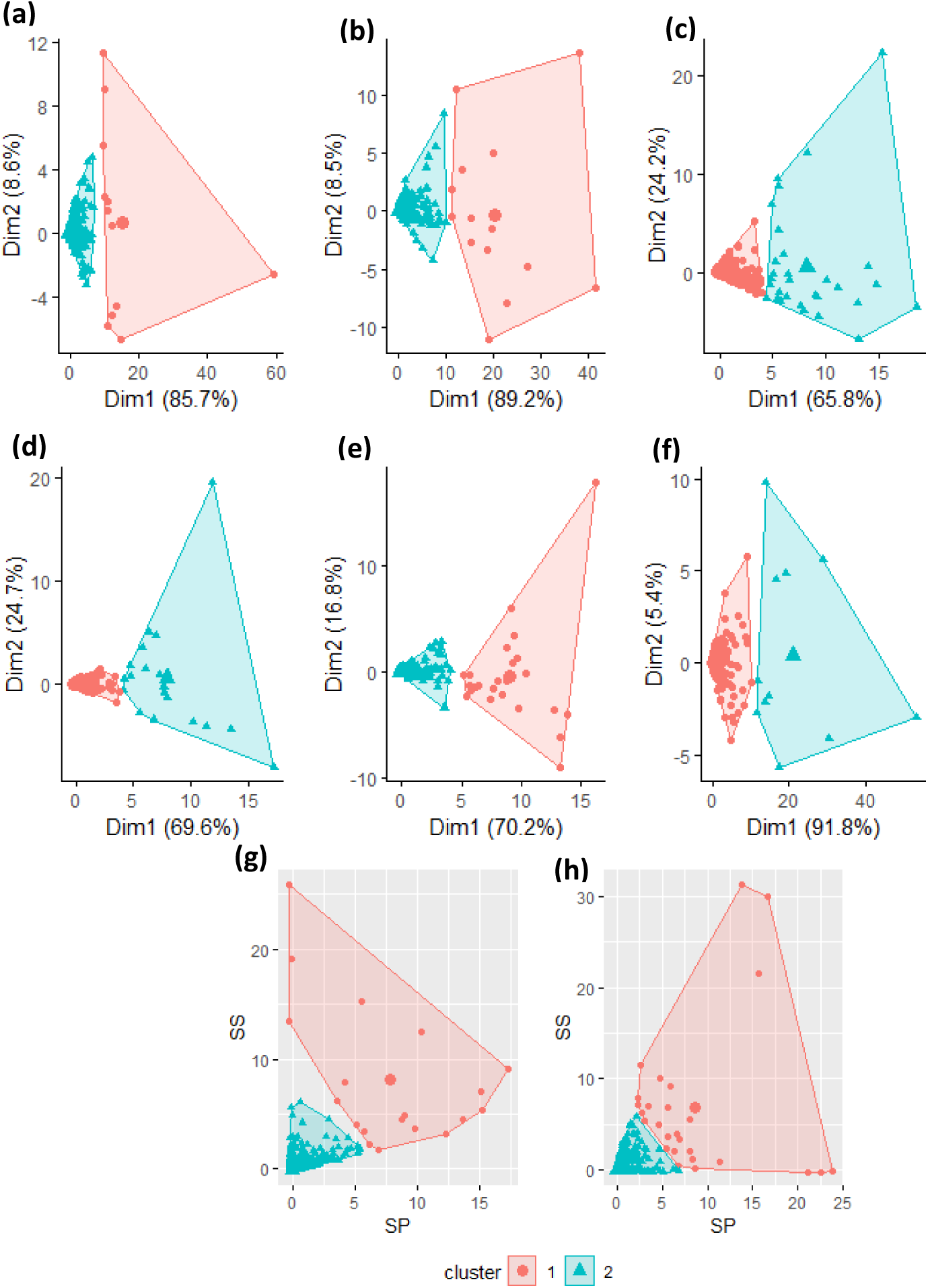
K-means clustering of differentially expressed transcripts between leaves and cotyledons of S. seuvioides and between C_3_ and C_4_ species. a: Up-regulated transcripts in leaves compared to CD (Up-L/L-vs-CD), b: Up-regulated transcripts in CD compared to leaves (Up-CD/L-vs-CD), c: Up-regulated transcripts in CN compared to CD (Up-CN/CN-vs-CD), d: Up-regulated transcripts in CD compared to CN (Up-CD/CN-vs-CD), e: Up-regulated transcripts in L compared to CN (Up-L/L-vs-CN), f: Up-regulated transcripts in CN compared to L (Up-CN/L-vsCN), g: Up-regulated transcripts in C_4_ compared to C_3_ (Up-C_4_/C_4_-vs-C_3_), h: Up-regulated transcripts in C_3_ compared to C_4_ (Up-C_3_/C_4_-vs-C_3)_. Colours represent clusters

**Fig. 9.**
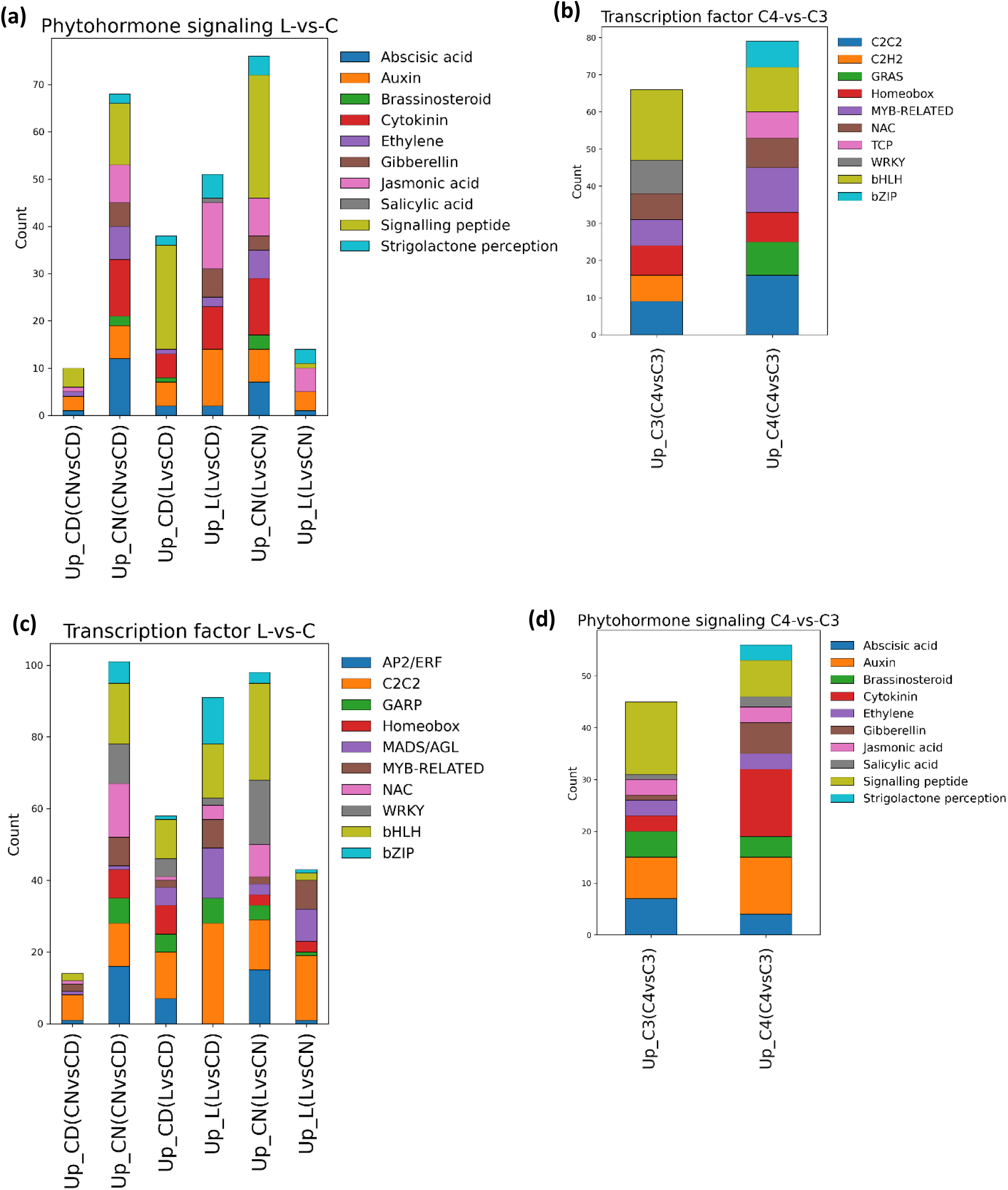
Stacked bar charts of functional annotated differential expressed genes and abundance of selected genes involved in CCM and starch metabolism. (a) Functional annotated differential expressed genes related to transcription factors between leaves (L) and cotyledons (C). (b) Functional annotated differential expressed genes related to transcription factors between C_3_ and C_4_ species. (c) Functional annotated differential expressed genes related to phytohormones between L and C. (d) Functional annotated differential expressed genes related to phytohormones between C_3_ and C_4_ species

Our results indicated that the same TF families are recruited to a variable degree for controlling cotyledons and leaves of *S. sesuvioides*. The top 10 TF families abundant in cotyledons and leaves during the day and in CN were basic Helix-Loop-Helix (bHLH), C2C2, APETALA2/ethylene-responsive factor (AP2/ERF), WRKY, MADS/AGL, bZIP (Basic Leucine zipper), NAC, Myoloblastosis (MYB)-related, GARP (Golden2, ARR-B, Psr1), and Homeobox (Fig. 9a). While during the day, the number of TFs related to C2C2, BHLH, and bZIP families were higher in leaves than in cotyledons, no TFs related to Homeobox and AP2/ERF were enriched in leaves. When comparing CD and CN, we found nearly all TF families were abundant at night. We then specifically looked at TFs that are frequently expressed in these families. Genes were selected by their potential involvement in the regulation of CAM and C_4_ photosynthesis. Our data showed the TFs bHLH87, REVEILLE1, WRKY40, NAC83, and SCL15 were up-regulated in the cotyledons during the night, whereas TFs BBX24, and SCL14 were significantly abundant in the leaves (Fig. 4c). To confirm whether these TFs regulate C_4_ and CAM photosynthesis, we explored TF binding sites in sequences of transcripts related to C_4_ photosynthesis and starch metabolism (Table 2). Motif enrichment analysis revealed elements AAAAAG and CTTTTT from the C2C2-DOF family were the most enriched in leaves during the day, while element CTTTTT was the most enriched in cotyledons (Table 2). When comparing CD and CN, element CGCCGCC from the AP2/ERF family was enriched in CD whereas element (CTTTTT) was enriched in all comparisons.

**Table 2:**
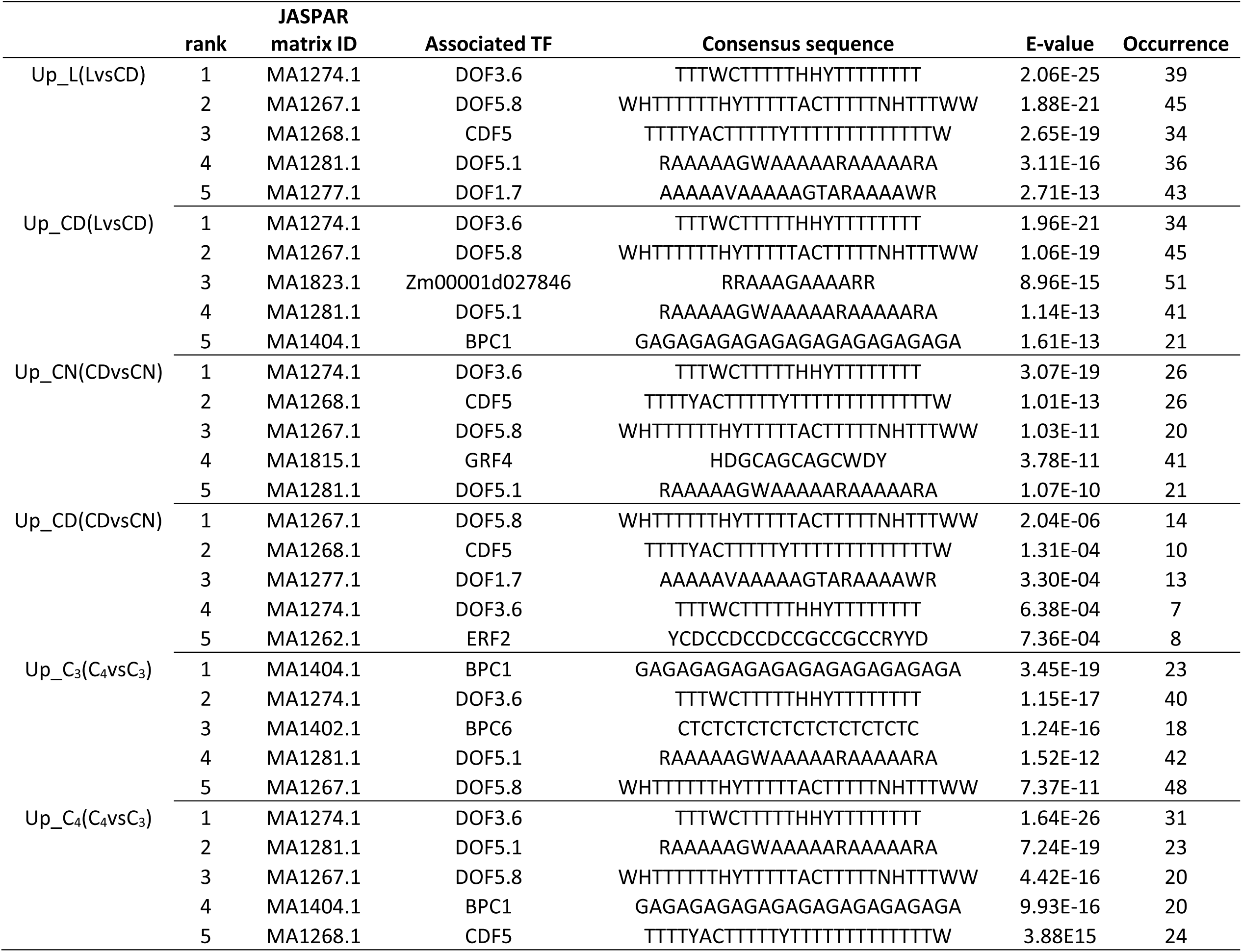
Top five motif enrichments of differentially expressed genes related to CCM (Carbon Concentrating Mechanism) and starch metabolism in *Sesuvium* species.

For comparison between *S. sesuvioides* and *S. portulacastrum*, TF families C2C2, C2H2, GRAS, Homeobox, MYB-RELATED, NAC, Teosinte branched1/cycloidea/proliferating cell factor (TCP), WRKY, bHLH, and bZIP were found on top of the list (Fig. 9b). Except for TFs from the GRAS family present only in C_4_-like, all others were found in C_4_-like and C_3_ species (Supplementary Dataset S3-S4). We selected based on the literature candidate TFs that are potentially involved in the regulation of C_4_ and CAM (Fig. 4g). Different isoforms are recruited in the regulation of C_3_ and C_4_ species. Our data showed that REVEILLE 8, BHLH143, SCL6, COL5, and DOF1.2 were significantly up-regulated in C_4_ species, whereas REVEILLE 6, COL15, NFYA7, and HB6 were up-regulated in the C_3_ species (Fig. 4g, Supplementary Dataset S3-S4). We identified 63 nonredundant motifs (p < 0.01, E-value < 0.05) in sequences of genes upregulated in the C_4_-like species while 36 motifs were found in the C_3_ species (Supplementary Dataset S8. The top five of the most enriched motifs were annotated to C2C2-DOF families in C_4_-like and the C_3_ species with the element CTTTTT (Table 2). Although the most enriched motifs were similar between the C_4_-like species and the C_3_ species, the second most enriched motifs were elements (GAGA, BBR/BPC family) and (CACCAACM, MYB family) in the C_4_-like and C_3_ species, respectively. We found several motifs often present in the same transcript sequences. This suggests a coordinated and regulatory network of TFs controlling photosynthesis in which motif CTTTTT is dominant.

Phytohormones play critical roles in photosynthesis regulation and developmental processes ranging from organ initiation to senescence (Müller and Munne-Bosch, 2021). Moreover, phytohormones have been shown to mediate TF action in C_4_ and CAM plants (Ferrari et al., 2022). When comparing leaves to cotyledons, signaling peptides, auxin, cytokinin, jasmonic, and abscisic acids were the most abundant signaling hormones (Fig. 9c). We found that the expression of CYP707A4 (abscisic acid), LHW (cytokinin), and JAR6 (jasmonic acid) increased significantly in abundance in cotyledons at night (Fig. 4d). In cotyledons during the day, MS17 (Auxin) and GAST1 (signaling peptide) were up-regulated, while IAA14 (auxin) and GASA1 (signaling peptide) were significantly accumulated in the leaves. Many other signaling protein genes were found specific to leaves and cotyledons and are listed in (Supplementary Tables S6, S7). When comparing *S. sesuvioides* to *S. portulacastrum*, we found that signaling peptides, auxin, cytokinin, abscisic acid, and brassinosteroid were the top five endogenous signaling hormones (Fig. 9d). There were many genes related to hormonal signals that were specific to either C_3_ species or C_4_ species (Supplementary Tables S3, S4). For instance, genes PHP2 (cytokinin), RALF4 (signaling peptide), and IAA9 (auxin) were significantly accumulated in the C_3_ species while EIR3 (auxin), ERF118 (cytokinin) and CEPR2 (signaling peptide) were significantly accumulated in the C_4_-like species (Fig. 4h).

## DISCUSSION

### Photosynthetic mode in *Sesuvium* species

Our findings confirmed that adult leaves of *S. sesuvioides* perform C_4_-like photosynthesis with core C_4_ enzymes up-regulated. These up-regulated enzymes were involved in carboxylation (βCA, PEPC1), acid regeneration (ALAAT2, AspAT, NADP-MDH), decarboyxlation (NADP-ME, NAD-ME), transporters (BASS2, DIT1, DIT2, PPT2) and PEP regeneration (PPDK). This result is in accordance with anatomical, biochemical, and physiological observations (Bohley et al., 2019). The C_4_-like status of *S. sesuvioides* becomes evident in the still relatively high expression of photorespiratory genes (Fig. 3c).

Intriguingly, when comparing leaves of the adult plants with cotyledons of *S. sesuvioides*, the main carboxylation enzyme was PEPC isoform 4 (PEPC4) in leaves. There are several plant PEPC copies that are classified into photosynthetic (C_4_ and CAM) and non-photosynthetic isoforms (Leary et al., 2011). The PEPC4 isoform identified and up-regulated in the leaves is homologous to Arabidopsis AtPEPC4 which is involved in photosynthesis and may indicate a similar role in *S. sesuvioides*. In line with this result, PEPC3 and PEPC4 were significantly expressed in adult C_3_ leaves of *S. portulacastrum* when compared to the leaves of C_4_-like *S. sesuvioides*. This could also be explained by the fact that the C_3_ species *S. portulacastrum* induces weak CAM photosynthesis under drought stress (Winter et al., 2019). Heat and drought stresses repress nitrogen metabolism enzymes (Xia et al., 2020). Thus, regulation of nitrogen metabolism is crucial to maintain plant growth under stress conditions. PEPC plays a crucial role in C_4_ photosynthesis and in modulating the balance of carbon and nitrogen metabolism in *Arabidopsis* (Shi et al., 2015). This indicates that the up-regulation of these PEPC copies may be important to maintain the photosynthetic system and plant growth of *Sesuvium* species under adverse conditions. The biological role and localization of these PEPC copies in *Sesuvium* need to be further investigated, however, our results indicate that different PEPC copies are optimized for different photosynthetic functions in leaves and cotyledons of *Sesuvium*.

Generally, there are three subtypes of C_4_ photosynthesis depending on the decarboxylation enzymes. The main decarboxylation enzyme in *S. sesuvioides* was NADP-ME. The up-regulation of NADP-ME indicates that *S. sesuvioides* is employing NADP-ME in its decarboxylation mechanism. This result is consistent with biochemical and physiological observations in *S. sesuvioides* (Bohley et al., 2019). Moreover, transcriptome comparison between leaves of *S. sesuvioides* and *S. portulacastrum* revealed that *S. sesuvioides* can additionally or occasionally employ NAD-ME as decarboxylation enzymes (Supplementary Table S3).

### Photosynthetic plasticity in *Sesuvium*

Plants exhibit plasticity for a wide variety of ecologically important traits to adjust to environmental changes (Sultan, 2000). Photosynthetic plasticity underpins the ability of plants to acclimate and grow in adverse environments and may depend on plant ontogeny. Our data provide evidence of photosynthetic plasticity in *S. sesuvioides* (Aizoaceae) with C_4_ and CAM photosynthesis in leaves and cotyledons, respectively. During the day, decarboxylating enzymes were more strongly expressed in cotyledons compared to leaves while carboxylating enzymes were strongly expressed in leaves. This result was further confirmed by titratable acidity, which showed a significant accumulation of acids overnight in cotyledons. The co-occurrence of C_4_ and CAM photosynthesis has already been reported in Aizoaceae, namely for *Trianthema portulacastrum* (Winter et al., 2021), but this is the first time to report ontogenetic variability with respect to photosynthesis in the Aizoaceae family with CAM in cotyledons and C_4_ in leaves. In Amaranthaceae (incl. Chenopodiaceae), Lauterbach et al. (2017) based on RNA expression profiles showed the transition from C_3_ photosynthesis in cotyledons to C_4_ photosynthesis in adult leaves of *Salsola soda*. This phenomenon seems to occur in several species of Salsoleae according to C_3_-like features such as lower carbon isotope ratios and lack of Kranz anatomy in cotyledons (e.g., Pyankov et al., 1999; Pyankov et al., 2000). However, these species have never been tested for CAM metabolism. The presence of CAM in cotyledons may be induced by environmental clues. Indeed, no CAM was observed in the cotyledons and leaves of *S. sesuvioides* under well-watered conditions (Bohley et al., 2019). In the climate chamber, stressful conditions were mainly created by the maximum light intensity. CAM induction has been linked to a photoprotective role in *Portulaca oleracea* (Ferrari et al., 2020). This suggests a photoprotective role of CAM induction in cotyledons of *S. sesuvioides*.

### Integration of C_4_ and CAM photosynthesis

Our data suggested a possible co-occurrence of C_4_ and CAM photosynthesis in a single leaf of *S. sesuvioides* under adverse conditions (Fig. 5, Fig. 7a). In CAM photosynthesis, nocturnally accumulated malate is translocated out of the vacuole by a malate channel for subsequent decarboxylation during the light period. In *P. oleracea*, a C_4_ species that performs CAM when drought-stressed (Moreno-Villena et al., 2022), AtALMT9 that has been associated with CAM function (Ferrari et al. 2020) is a vacuolar malate channel (Kovermann et al. 2007). Interestingly, we found ALMT9 was significantly abundant during the light period in *S. sesuvioides*. Thus, the up-regulation of ALMT9 in leaves of *S. sesuvioides*, suggests that ALMT9 may function as a vacuolar malate efflux channel in *S. sesuvioides* and is therefore linked to CAM function. Taking all results together, this implies the integration of the hybrid system C_4_+CAM in *S. sesuvioides* under stress conditions. However, the modularity of this integration needs to be investigated. This co-occurrence of C_4_ and CAM in a single leaf in *S. sesuvioides* is probably facilitated by the particular C_4_-like phenotype of *S. sesuvioides* leaves where Rubisco is present in the mesophyll cells. The mesophyll cells of *S. sesuvioides* are succulent and outnumber the Kranz cells by two-fold. When the leaves grow older, the mesophyll portion becomes even larger and the carbon isotope ratios drop (Bohley et al. 2019). This might indicate a photosynthetic plasticity towards a higher proportion of CAM and or C_3_ relative to C_4_ in older leaves depending on the growing conditions.

### Regulation of photosynthesis and hormonal signaling in *Sesuvium*

The ability of plants to choose between photosynthetic pathways is controlled by TFs. Six TF families i.e., C2C2, Homeobox, NAC, WRKY, bHLH, and bZIP were found up-regulated in leaves and cotyledons of *S. sesuvioides* when compared leaves to cotyledons and also when compared to adult leaves of *S*. *sesuvioides* and to adult leaves of *S. portulacastrum*. These families have been hypothesized to be involved in the regulation of C_4_ and CAM photosynthesis in Chenopodiaceae, Aizoaceae, and Asteraceae (Siadjeu et al., 2021, Ferrari et al., 2022). However, TFs from the C2C2, and bHLH families were the most expressed in leaves, and cotyledons during day and night. Likewise, these TFs were predominant in the C_3_ and C_4_-like species. This indicates the significant weight of the C2C2 and bHLH TF families in the regulation of cotyledons and leaf development in *S. sesuvioides* and *S. portulacastrum*. The binding site of TF C_4_ zinc finger-type (DOF3.6) from the DOF/C2H2 was most enriched in C_4_ and CAM genes in all comparisons. In addition, at least three copies of DOFs were among the top five. Several copies of DOF proteins (DOF1 and DOF2) were found involved in the regulation of the light-dependent C_4_ gene PEPC in maize with antagonist effects. While DOF1 activates C_4_ genes, DOF2 can activate or repress them (Yanagisawa and Sheen, 1998). These results indicate that different copies of DOF genes are likely involved in the regulation of C_4_ and CAM genes in *S. sesuvioides* and *S. portulacastrum*, as well.

Transcription factors are regulated by phytohormones (signaling molecules) under environmental stresses (Saibo et al., 2009). Our data showed that phytohormones were clustered with C_4_ and CAM genes, as well as TFs which indicates a regulatory network involving TFs and phytohormones. Indeed, Ferrari et al. (2022) found that ABA and CK-related genes regulate TFs connected to CAM and C_4_ photosynthesis in *Portulaca oleracea*. While ABA and CKs have been studied intensely in CAM and C_4_ photosynthesis, several other phytohormones that regulate photosynthesis (reviewed by Müller and Munne-Bosch, 2021) have received little attention. Here, we found that transcripts encoding signaling peptides were the most abundant plant hormones during the day and at night in cotyledons as opposed to leaves. Similarly, diurnal and nocturnal comparison expression in cotyledons showed that signaling peptides were predominantly accumulated. Transcripts of genes encoding for the signaling peptides Gibberellic acid (GA)-Stimulated Arabidopsis/GA-Stimulated Transcript (GAST) and the Rapid Alkalinization Factor (RAFL) were predominantly expressed in cotyledons as compared to leaves. These plant hormones play important roles in plant growth, development, and stress responses (Wu et al., 2020; Lee et al., 2023), and may control cotyledon growth and response to environmental conditions in *S. sesuvioides*. Conversely, in leaves as opposed to cotyledons during the day, the most dominant hormone was jasmonic acid followed by auxin and cytokinin CKs. Jasmonic acid, auxin, and cytokinin are classical phytohormones that regulate various aspects of plant growth and abiotic and biotic stress responses. While jasmonic acid can regulate stomatal closure and opening under drought stress in *Arabidopsis* (Savchenko et al., 2014), auxin coordinates cell division, expansion, and differentiation (Perrot-Rechenmann, 2010), and CKs are implicated in cell cycle progression (Park et al., 2021). Several genes encoding for these phytohormones were found to be significantly accumulated (Supplementary Tables S6, S7) and should be used as candidate genes involved in the regulation of CCMs in these species.

## Conclusions

This study of gene expression profiles of *S. sesuvioides* provides evidence of extraordinary photosynthetic plasticity under adverse conditions with induced CAM in cotyledons and an integration of CAM and C_4_-like photosynthesis in adult leaves. However, the modularity of the co-occurrence of the two CCMs needs to be explored in future studies. We assume that further detection of co-occurring CCMs is just a matter of more experimental studies that explicitly look for this and we believe that it is more common in succulent C_4_ lineages than currently known. Our findings suggest a complex regulatory network involving TFs and phytohormones and underpin the regulation of CCMs and adaptation of *Sesuvium* species which grow in disturbed and highly dynamic environments.

## Supplementary data

**Fig. S1:** Images of potted *S. sesuvioides* plants growing in the climate chamber and in the uncontrolled greenhouse environment.

**Fig. S2** The best K values were determined using the Elbow and Silhouette methods, as well as the Gap Statistic, in all comparisons.

**Dataset S1** Up-regulated transcripts, count, and annotation in CD as compared to CN of *S. sesuvioides*

**Dataset S2** Up-regulated transcripts, count, and annotation in CN the day as compared to CD of *S. sesuvioides*

**Dataset S3** Up-regulated transcripts, count, and annotation in *S. sesuvioides* compared to *S. portulacastrum*

**Dataset S4** Up-regulated transcripts, count, and annotation in *S. portulacastrum* as compared to *S. portulacastrum*

**Dataset S5:** List of differentially expressed transcripts within clusters in all comparisons

**Dataset S6** Up-regulated transcripts, count, and annotation in leaves compared to cotyledons of *S. sesuvioides*

**Dataset S7** Up-regulated transcripts, count, and annotation in cotyledons compared to leaves of *S. sesuvioides*

## Acknowledgments

We would like to thank Sebastian Walter for germinating the seeds of *S. sesuvioides* and growing the plant material. Additionally, we extend our thanks to Thibaud Messerschmid for reviewing and providing feedback on the initial manuscript.

## Author contributions

Conceptualization: GK; Data curation: CS; Formal Analysis: CS; Funding acquisition: GK; Investigation: CS; Methodology: CS; Project administration: GK; Resources: GK; Software: CS; Supervision: GK; Validation: CS, GK; Visualization: CS; Writing-original draft: CS; Writing – review & editing: GK, CS.

## Conflict of interests

The authors declare no competing interests.

## Funding

This work was supported by the Deutsche Forschungsgemeinschaft (DFG) with grants to GK (KA1816/7-3).

## Data Availability

## Notes

### Competing Interest Statement

The authors have declared no competing interest.

